# Influence of antibody-antigen interaction on fluorescence of Rhodamine 6G conjugated to the antibodies measured in human serum

**DOI:** 10.64898/2026.11.03.763607

**Authors:** Dinesh Thiyagaraj, Alessandro Del Re, Quan Duc Pham, Irene Gómez Garrote, Ahmad Saudi, Oleh Fedorych

## Abstract

Fluorescent dyes conjugated to monoclonal antibodies specifically trastuzumab (TRAS) and pertuzumab (PERT) serve as sensitive reporters of changes in the local energetic environment arising from antibody-antibody and antibody-antigen interactions. Upon covalent conjugation, the antibody local environment perturbates the energetic state of the dye, resulting in changes of the fluorescence emission energy and emission width of the dye. The dependence of these spectral parameters on antibody concentration was measured in HBS-8, diluted and undiluted human serum. In addition, the emission energy and emission width of dye-conjugated TRAS were investigated as a function of recombinant human epidermal growth factor receptor 2 (HER2) extracellular domain concentration in HBS-8 and in diluted and undiluted human serum. Antibody-antibody and antibody-antigen interactions produce remarkable contributions into the emission energy and emission width. These contributions depend on the composition of the surrounding medium. Our study showed that the samples containing 100 *nM* TRAS in serum exhibited the largest spectral changes upon interaction with HER2 suggesting that antibody-antigen interaction are not linear across concentration range of the antibodies. Moreover, our results demonstrate that interactions between dye-conjugated antibodies and their binding partners can be detected under physiologically relevant conditions. These findings establish fluorescence emission energy and emission width analysis as a potential approach for probing antibody interactions in complex biological matrices.

## Introduction

Monoclonal antibodies play an increasingly important role in modern therapeutics, creating a growing need for experimental methods capable of characterizing their interactions with target molecules under biologically relevant conditions. Trastuzumab (TRAS) and pertuzumab (PERT) are the most used therapeutic monoclonal antibodies targeting human epidermal growth factor receptor 2 (HER2) [1-4]. There are several established experimental techniques, including ELISA, SPR, and ITC [5], used to characterize antibody-antigen interactions. However, many of these methods require purified antibodies and strictly defined buffers. And quantitative determination of the interactions in therapeutically relevant fluids such as blood, plasma, or serum most likely will fail.

One of approaches for the antibody-antigen interaction analysis in therapeutically relevant fluids is labeling of antibodies through covalent conjugation with fluorescent dyes [6-8]. Although conjugation requires chemical modification of the antibodies and may potentially alter some of its physicochemical properties, it provides a highly selective optical probe that can be detected even in complex biological environments.

In the present work, fluorescent dyes are employed, not as light reporters but as energy sensors. Through covalent attachment to an antibody, the dye becomes sensitive to changes in the local energetic environment of the conjugated molecule. Interactions between an antibody and neighboring antibodies, proteins, or antigens modify this energetic environment and can be detected through changes in the fluorescence emission energy and emission width of the conjugated dye. In addition, it will be demonstrated that the environment, in this specific case, human serum has dramatic influence on the antibodies and their ability to interact with the binding partner. Moreover, characterization of antibody interactions in media different from the final biological environment leading to ambiguous conclusions can be solved with the disclosed approach.

## Materials and methods

Rhodamine 6G (Rh6G) 5-isomer was conjugated to TRAS and PERT at lysine residues using amine coupling chemistry. Trastuzumab (cat# 68046-H002), pertuzumab (cat# 68046-H001), and recombinant HER2 extracellular domain (cat# 10004-H08H) were purchased from Sino Biological. TRAS and PERT at 30 µ*M* were incubated with 100 µ*M* Rh6G NHS ester in a final volume of 100 µ*L* for 15 minutes at room temperature in the dark. Free unreacted Rh6G was removed using Zeba Dye and Biotin Removal Spin Columns (cat# A44296, Thermo Fisher Scientific) according to the manufacturer’s protocol. Each spectroscopic measurement was performed on a 100 µ*L* sample at room temperature. The interaction of Rh6G-conjugated trastuzumab and pertuzumab with recombinant HER2 extracellular domain was studied at concentrations up to 350 *nM* in assay buffer HBS-8 (100 *mM* HEPES, pH 8.0, 150 *mM* NaCl) and in different dilutions of normal human serum (11: 50, 1: 25, 1: 10, and undiluted).Human serum was purchased from Sigma-Aldrich (cat# P2918). Serum dilutions were made using PBS (137 *mM* NaCl, 2.7 *mM* KCl, 10 *mM*, Na_2_HPO_4_, 1.8 *mM* KH_2_PO_4_, pH7.4). To each 100 µ*L* antibody solution, 0.5 µ*L* of HER2 from the appropriate stock was added to achieve the desired final concentration.

### Theory and Model

It is well known that conjugation of fluorescent dyes to proteins can alter their optical properties [9, 10]. The changes are often discussed in terms of modifications to the dye’s local microenvironment [11, 12]. However, these discussions neglect quantitative measure of the influence exerted by the conjugated protein on the dye. In the present work, we will determine whether changes in fluorescence emission energy and emission width can be used to characterize the strength of intermolecular interactions involving an antibody and its antigen.

From quantum-mechanical perspective, conjugation of a dye to a macromolecule can be considered as a perturbation of its local environment which also can be described in terms of perturbation theory [13]. The perturbation on the dye emission arising from its conjugation with the macromolecule can be determined and the concept is explained in detail below.

On the S1 Fig. a) and b) shows the emission energy and spectral width of free Rh6G in HBS-8 and human serum, with the addition of TRAS antibody and HER2 antigen. Presence of TRAS or HER2 does not cause any remarkable changes in fluorescence of free Rh6G. The changes in the medium modify the fluorescence spectra the most. In HBS-8 and undiluted serum the emission energies are 2.2964 *eV* and 2.2884 *eV* and emission widths of 0.1074 *eV* and 0.1054 *eV*, respectively. As it will be shown, the emission energy and emission width of Rh6G conjugated with TRAS antibody are 2.3105 *eV* and 0.145 *eV* in HBS-8, and 2.301 *eV* and 0.145 *eV* in serum.

The difference between free Rh6G and TRAS conjugated with Rh6G shall be used for estimation of the effective perturbation potential, see Supplementary Information section. The estimated value of the effective perturbation potential is around 100 *meV* (98.4 *meV* and 100.4 *meV* in HBS-8 and serum, respectively). This value substantially exceeds the thermal energy (~25 *meV*), showing that the perturbation induced by the macromolecule is energetically significant and the dye may serve as a sensitive reporter of energetic rearrangements occurring with the antibody.

Fig.1 presents the energy diagram demonstrating how conjugation of the dye (Rh6G) with the antibody (TRAS) via an amino linkage changes the dye emission. Initially, the emission properties of free Rh6G are determined by the surrounding medium and are characterized by an emission energy *E*_*D*_ and emission width σ_*D*_. Covalent linkage of the dye to the antibody creates a perturbation potential *U* magnitude of which increases emission energy of the conjugate to *E*_*C*_ and broadens the emission to σ_*C*_. Interaction with another antibody may rearrange the strength of perturbation potential to a *U*^′^ so the dye will change it emission energy and emission width as 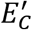 and 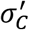 respectively. It also shows that *U*^′^ is dependent on mean distance between antibodies *r*. Adding HER2 protein to the antibody solution induces change of the perturbation potential (*U*^′′^) due to antibody-antigen interaction which results in the emission energy 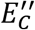 and emission width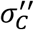. Finally, interaction between antibodies and antigens (all with all) induces perturbation potential *U*^′′′^ which cause the emission energy 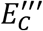 and emission width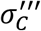.

As it will be shown later, the perturbation potential *U* is influenced by the interaction of the antibody with another antibody or protein or antigen, and this influence is reflected in the dye’s emission. The main challenge resolved in the research is determination of effect arising from this interaction. Since the interaction between the antibody and one of its interaction partners is dependent on the intermolecular distance *r* between them. Therefore, the concentration-dependent measurements will provide explicit insight into the interaction.

Following the exchange-narrowing theory [14-17], where the interaction-induced perturbation is assumed to be governed by an interaction term *κ*exp(− *r*/λ). The spectral response was modified and represented as

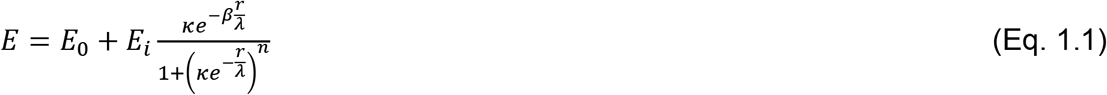

And

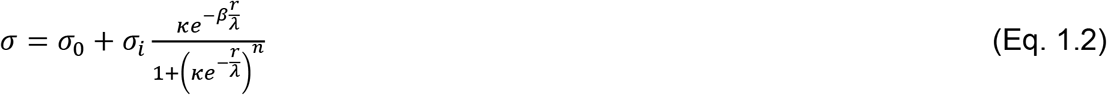

where *E*_0_ and σ_0_ are the emission energy and emission width in the absence of interaction. Depending on the molecular configuration *E*_0_ and σ_0_ can be represented by 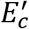 and 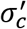 (Ab-Ab configuration) or 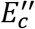 and 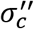 (Ab-Ag configuration) or 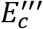 and 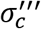 (Ab-Ab, Ab-Ag, Ag-Ag). The interaction-induced emission energy shift and emission width variation are described as *E*_*i*_ and σ_*i*_. Interaction induced contributions appearing from Ab-Ab interaction are 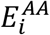 and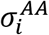, from Ab-Ag interaction are 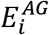 and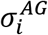. Interaction terms induced by all contributions (Ab-Ab, Ab-Ag, Ag-Ag) are 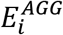 and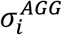. The interaction amplitude is given by *κ*, λ is the interaction transfer length, *n*controls the transition between the exchange-narrowing and exchange-broadening regimes. Accordingly, with [15] *n*= 2, but to reflect that the interaction involves multiple molecular species and multiple interactions, *n*becomes larger than 2. The parameter *β* corresponds to an effective interaction transfer parameter. It was introduced into exchange-narrowing theory to allow us to reflect the efficiency of exchange interaction spreading by the surrounding. It emphasizes that in buffers without proteins the exchange interactions decaying with distance more rapidly than in buffers containing serum.

### Experiment and Discussion

Despite the considerable interest in antibody-antibody and antibody-antigen interactions in complex media, studies of such interactions in media other than highly purified buffers appear to be largely absent. In this research, we performed a range of experiments in which interactions between antibodies and antigens were measured in HBS-8 and in diluted and undiluted serum. The focus was on the anti-HER2 antibody TRAS and a recombinant HER2 protein. Antibody-antibody interactions in serum and its dilutions were also investigated using the anti-HER2 antibody PERT.

Details of the experimental technique can be found elsewhere [18,19]. The signal we measured originated from the dye-conjugated antibody. Before using complex media and analyzing complex interactions, we investigated how the antibody emission energy and emission width changes with its concentration increase in HBS-8. The TRAS emission energy and emission width for increasing TRAS concentration, corresponding to decreasing TRAS-TRAS mean intermolecular distance, in HBS-8, is shown in Fig. 1a) and b). Both the emission energy and the emission width decrease with decreasing intermolecular distance. The reduction in energy and the narrowing of the emission width stabilize at around 200 *nM* (corresponding to an intermolecular distance of approximately 200 *nm*), and at higher concentrations (shorter distances) a slight increase in emission energy and broadening of the emission are observed. The red dashed line in both plots corresponds to the numerically obtained model. For the emission energy, 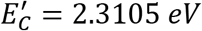, 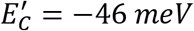, *κ* = 2.2, λ = 220 *nm, n*= 2, and *β* = 1. For the emission width, 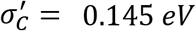 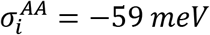, *κ* = 2.2, λ = 290 *nm, n*= 2, and *β* = 1.

**Fig. 1.**
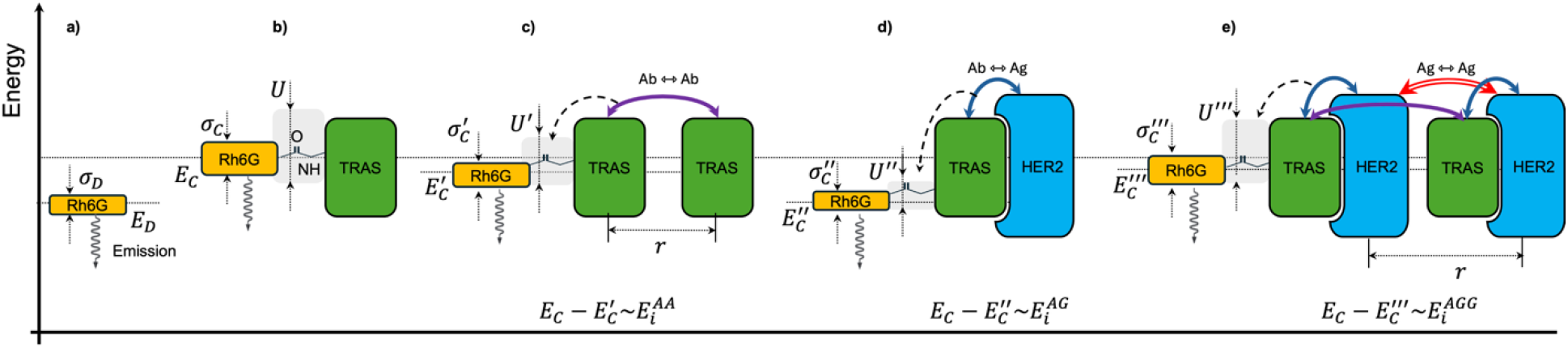
**a)** shows the energy diagram representing emission energy ***ED*** and emission width **σ*D*** of free Rh6G (orange rectangle). **b)** shows the emission energy ***EC*** and emission width **σ*C***of Rh6G in result of conjugation to TRAS (green rectangle) and the perturbation potential caused by conjugation is denoted as ***U*. c)** distance ***(r)***dependent antibody-antibody interaction (Ab⇿Ab) rearrangement the perturbation potential ***U*** as ***U***^′^ which it results in the emission energy 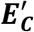 and emission width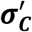. **d)** antibody-antigen interaction (Ab⇿Ag) rearranges the perturbation potential ***U*** as ***U***^′′^ and it results in the emission energy 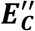and emission width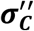. **e)** shows the rearrangement of the perturbation potential ***U*** as ***U***^′′′^, which includes Ab⇿Ab, Ab⇿Ag as well as distance ***(r)***dependent antigen-antigen (Ag⇿Ag) contributions, it results in the emission energy 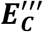 and emission width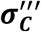.

The presence of serum induced an immediate response from the dye-conjugated TRAS antibodies. For the first serum dilution 1: 50, the emission energy shows a continuous increase with decreasing TRAS-TRAS mean intermolecular distance, as shown in Fig. 1c) and d). For distances shorter than 200 *nm*, a rapid increase is observed in both the emission energy and the emission width. For the 1: 25 serum dilution, the emission energy and emission width initially decrease and then increase again for distances shorter than 200 *nm*.

The obtained mean intermolecular distance dependences were described using a similar set of parameters. First, the 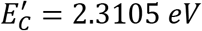 determined for HBS-8 changed to 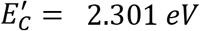 for all buffers comprising serum. The emission energy of the dye-conjugated TRAS antibody in the presence of serum is around 10 *meV* lower than in HBS-8. No change in emission width between buffers was registered. The emission width of the isolated dye-conjugated antibody remained the same in HBS-8 and in the serum-containing liquids,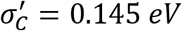.

The interaction profile of the antibody changes significantly in the presence of serum. The *n*parameter increases from *n*= 2 (without serum) to approximately *n*= 12 (with serum). This is especially pronounced for the 1: 50 and 1: 25 serum dilutions, where, at distances shorter than 200 *nm*, a sharp increase in the emission energy and emission width was observed. Another parameter affected by the presence of serum is *κ*, which changes from *κ* = 2.2 for HBS-8 to *κ* = 1.55 for serum and its dilutions. The amplitude of antibody-antibody interaction decreased, and the interaction transfer length remains largely conserved.

As seen for the 1: 50 serum dilution in Fig. 1c) and d), a decrease in the intermolecular distance initially increases the emission energy and broadens the emission width. To model this behavior, the interaction transfer parameter *β* assumed to be *β* = −0.07. At higher serum content, the interaction transfer efficiency increases, with *β* = 0.19, 0.4, and 0.4 for the 1: 25 Fig. 1e) and f), 1: 10 Fig. 1g) and h), and 1: 0 Fig. 1i) and j) dilutions, respectively, but it remains smaller than the *β* = 1 observed in HBS-8. Showing that in HBS-8 the antibody-antibody interaction decaying faster with decreasing mean antibody-antibody distance. The conservation of the interaction transfer length λ and reduction of the interaction amplitude *κ*, as well as the reduction of the interaction decay rate *β* suggest participation of another molecule which indirectly affecting on the antibody-antibody interactions. We believe that this molecule could be the proteins present in the serum.

Interestingly, despite the differences in the experimental conditions, measuring the antibody-antibody interaction in HBS-8 and serum using PERT gave results very similar to those obtained for TRAS Fig. 4a) and b). This suggests that the interaction-transfer mechanism is similar for TRAS and PERT antibodies.

The results for the TRAS-TRAS and PERT-PERT antibody interactions in serum and its dilutions are presented in Fig. 4. The emission energy change and the emission width variation for antibody-antibody interactions are shown in Fig. 4a). As it is seen, increasing the serum content, and consequently decreasing the distance between the protein molecules present in the serum, leads to a decrease in the interaction-induced emission energy and emission width.

This provides evidence for an indirect interaction between antibodies mediated by proteins present in serum. Comparison of the values obtained for antibody-antibody interactions in HBS-8 alone and in serum shows that the protein-mediated antibody-antibody interaction is at least four times weaker than the direct antibody-antibody interaction, with interaction-induced emission energies of 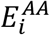 is equal to −11 *meV* and −46 *meV*, respectively. Similarly, the emission broadening in serum 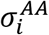 is approximately two times smaller than in HBS-8 alone, with values of −28 *meV* and −56 *meV*, respectively.

The magnitude of the perturbation potential rearrangement is reflected as shift of the emission energy. Stronger perturbation pushes the emission energy further from its free-dye value, whereas a reduction in the perturbation potential reduces the emission energy. When an antibody interacts with another molecule, the interaction modifies the energetic state of the coupled system and changes the resulting perturbation potential affecting on the dye, as shown in Fig. 1c). In other words, the larger the magnitude of 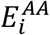 then stronger the interaction between the antibody and the molecule finally appears.

The parameters characterizing the spatial distribution of antibody-antibody interactions in the presence of the serum proteins are shown in Fig. 4b). Antibody-antibody interactions respond to the presence of the additional medium which transfers these interactions. Increasing the serum content increases interaction transfer effect, however it is not reaching the value obtained in HBS-8, where *β* = 1.

At protein concentrations higher than those present at the 1: 10 dilution have no further influence on the interaction transfer. The effective interaction distance also appears to reach a stationary value for serum concentrations above 1: 10. The analysis shows that the effective interaction distance is shifted toward smaller distances, corresponding to higher antibody concentrations, and is at least two times smaller than that determined for the HBS samples: approximately 200 *nm* in HBS-8 compared with approximately 220 *nm* in serum at 1: 10 dilution and in undiluted serum.

Next, we measured how the TRAS antibody binds to the recombinant HER2 protein. The measurements were performed in HBS-8, in different serum dilutions, and in undiluted serum. For this purpose, we selected three initial concentrations of the TRAS antibody, 30 *nM*, 100 *nM*, and 200 *nM*. These concentrations were chosen based on the TRAS-TRAS antibody interaction in HBS-8. As can be seen in Fig. 1a) and b), at 30 *nM* the antibodies are mainly isolated, and the antibody-antibody interaction is weak. At 100 *nM*, the antibodies are already coupled, while at 200 *nM* the antibody-antibody interaction is close to its absolute extremum.

Samples containing HBS-8 and the above-mentioned concentrations of dye-conjugated TRAS antibody were measured first. The recombinant HER2 protein was then added to the samples at a concentration of 10 *nM*, and the samples were measured again. This procedure was repeated several times, gradually increasing the concentration of the HER2 protein until a final concentration of 600 *nM* was reached.

Fig. 5a) and b) show the changes in emission energy and emission width of the TRAS antibody at the three initial antibody concentrations as the HER2 protein concentration increases, shown as the decrease of mean intermolecular distance between HER2 proteins. As can be seen, increasing the HER2 concentration increases the emission energy of the dye conjugated to TRAS. A similar trend is observed for the emission width. In contrast to the antibody-antibody interaction, where a reduction in emission energy was observed, TRAS-HER2 interaction results in an increase of the emission energy reported by the conjugated dye, indicating that HER2 modifies the perturbation potential acting on the fluorophore in a qualitatively different manner.

Binding of HER2 to the TRAS antibody modifies the perturbation potential *U*^′′′^, see Fig.1 e), therefore the emission energy of TRAS in HBS-8, 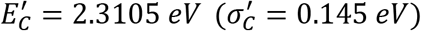, Fig. 2a), reduces to the emission energy 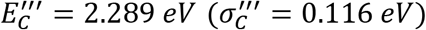, of the TRAS-HER2. TRAS-HER2 interaction induced emission energy shift is approximately 21.5 *meV* and emission width changed on 29 *meV* comparably to the TRAS without HER2 proteins. Moreover, the estimated magnitude of perturbation potential *U*^′′′^is around 44.4 *meV* which is still above thermal energy but twice lower than the perturbation caused by the covalent conjugation. The obtained result has particular importance since it shows that the dye conjugated to TRAS antibody can sense the antibody interaction with HER2.

**Fig. 2.**
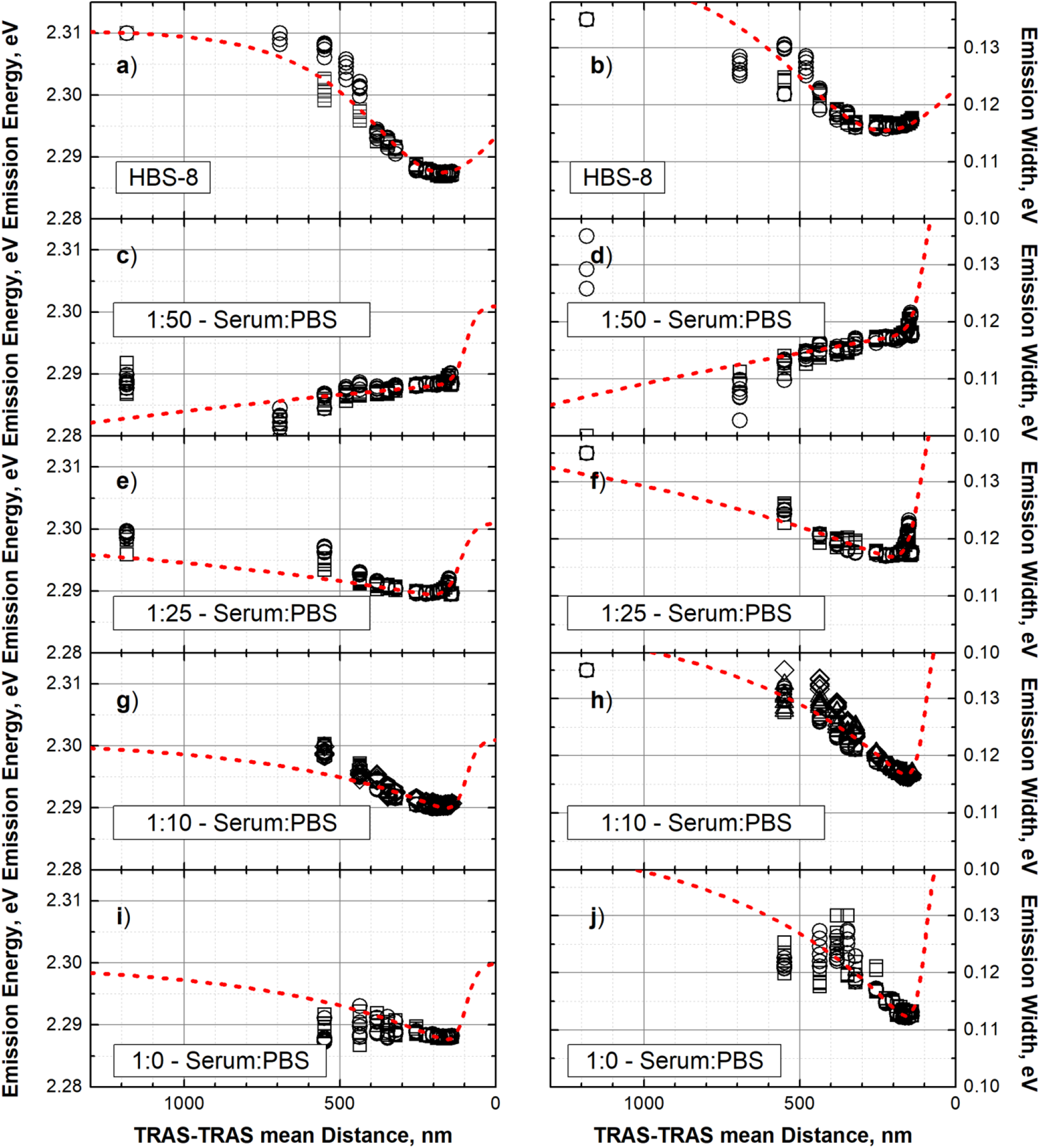
Dependence of emission energy **a), c), e), g), i)** and emission width **b), d), f), h), j)** of dye conjugated TRAS on TRAS-TRAS mean distance (concentration) measured in HBS-8, **1: 50, 1: 25, 1: 10** and **1: 0** serum. Experimental data are shown with the points, red dashed line corresponds to the theoretical model.

**Fig. 3.**
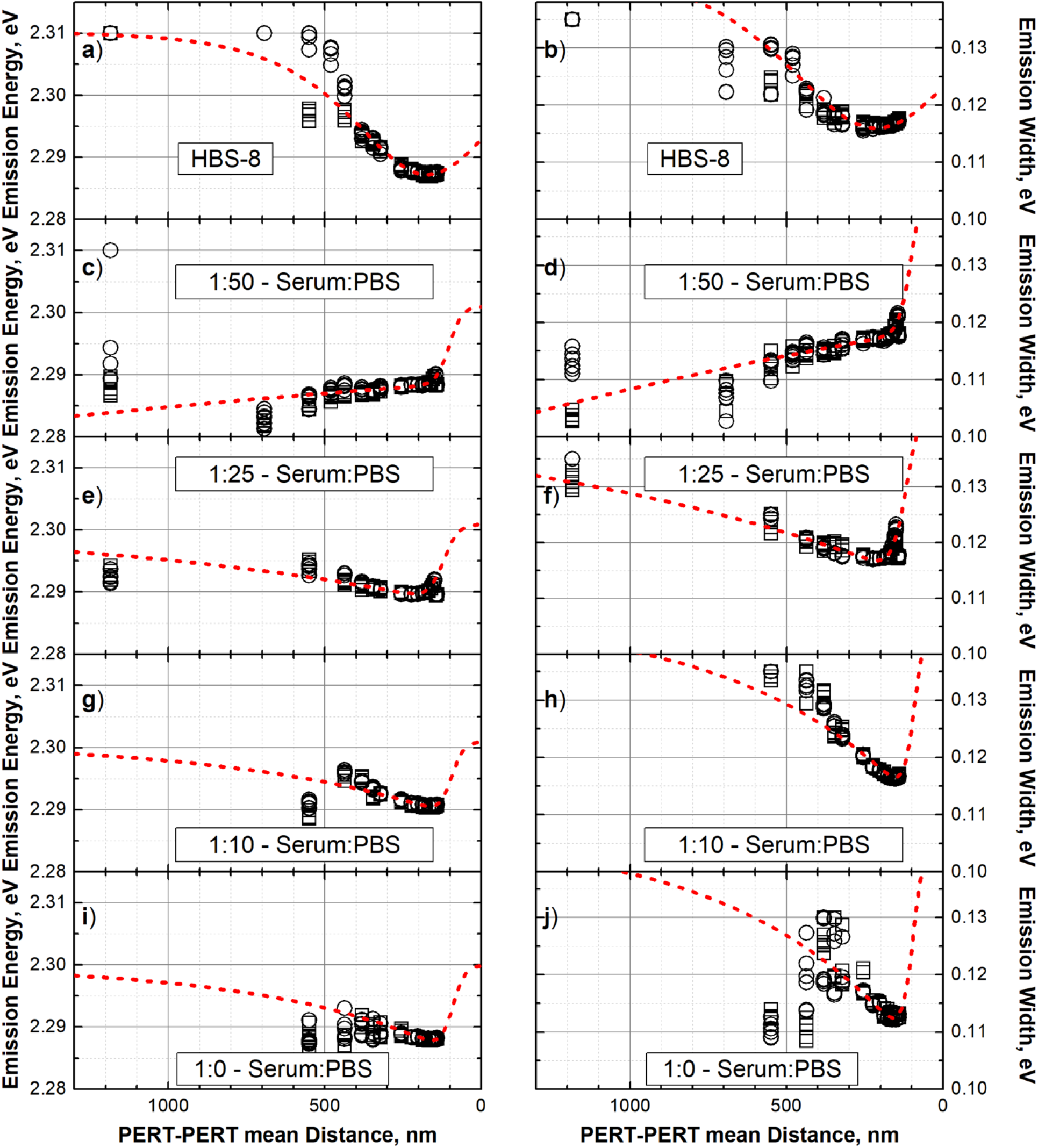
Dependence of emission energy **a), c), e), g), i)** and emission width **b), d), f), h), j)** of dye conjugated PERT on PERT-PERT mean distance (concentration) measured in HBS-8, **1: 50, 1: 25, 1: 10** and **1: 0** serum. Experimental data are shown with the points, red dashed line corresponds to the theoretical model.

**Fig. 4.**
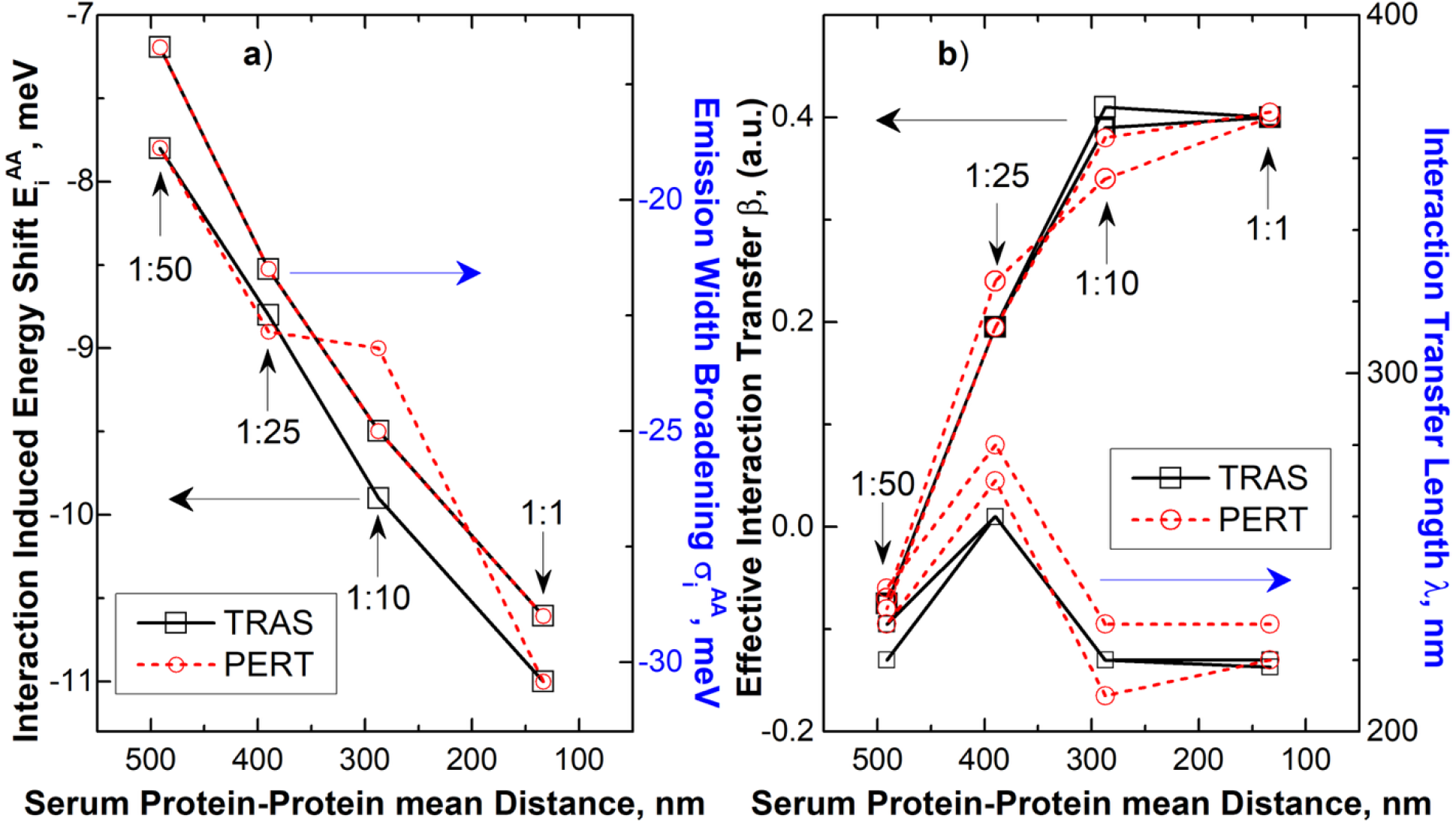
**a)** Interaction induced energy shift 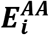 and emission width broadening 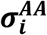 and **b)** effective interaction transfer ***β*** and interaction transfer length ***λ*** evaluated for TRAS (black squares with black solid line) and PERT (red points with red dashed line) in serum of different dilutions shown as serum protein-protein mean distance.

**Fig. 5.**
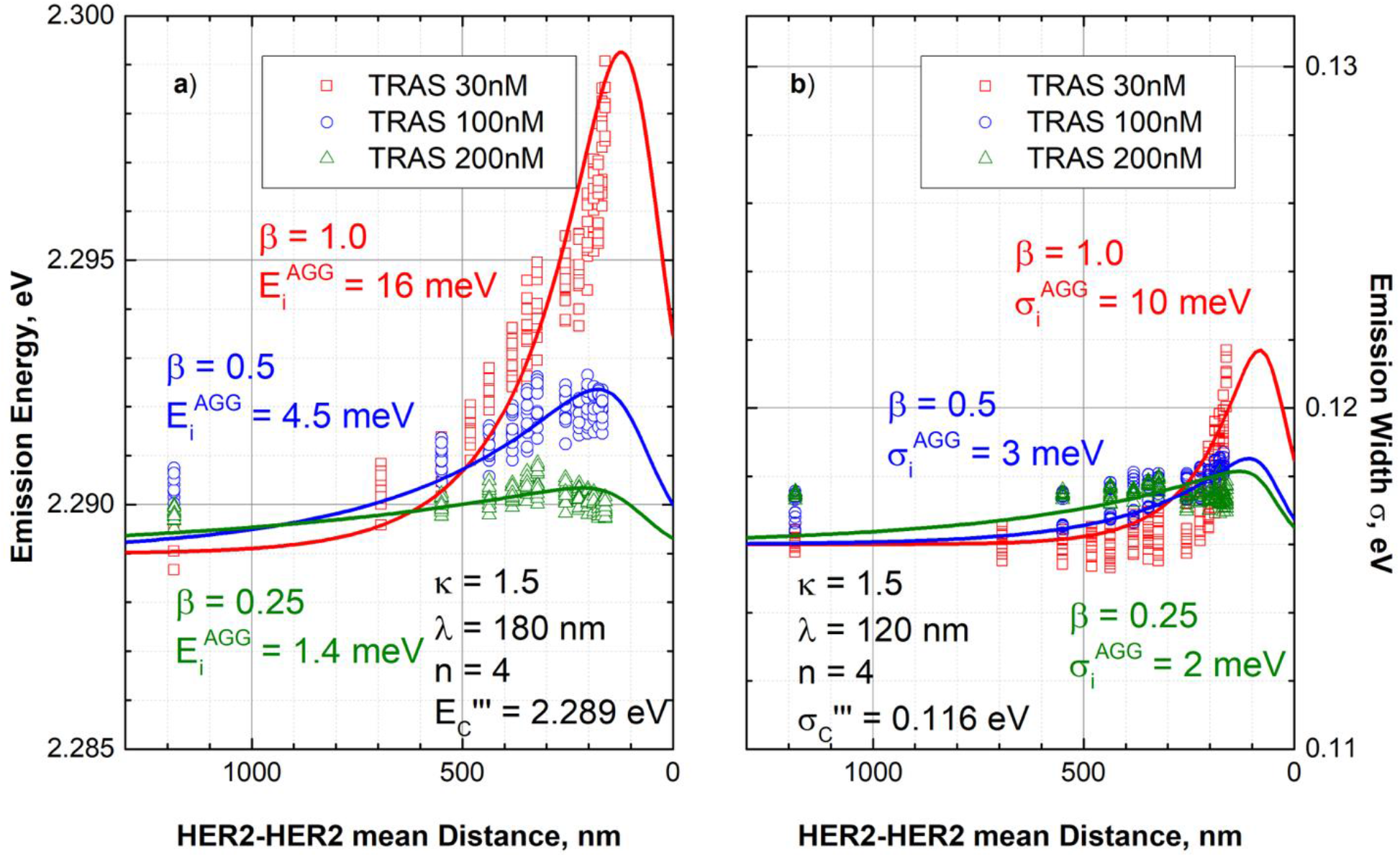
Dependence of TRAS **30 *nM*** (red squares), TRAS 100 ***nM*** (blue points) and TRAS **200 *nM*** (green triangles) emission energy a) and emission width b) on HER2 concentration presented as HER2-HER2 mean distance in HBS-8. Theoretically obtained model curves are shown with solid lines, **30 *nM*** solid red line, **100 *nM*** solid blue line and **200 *nM*** solid green line.

For all three TRAS concentrations after adding of 10 *nM* of HER2 the measured values of emission energy and emission width are the same 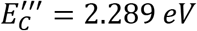 and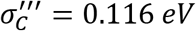, respectively. However, the TRAS only samples showed emission energy described as 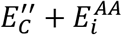 and emission width described as 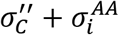 equal to 2.3013 *eV* and 0.124 *eV* for 30 *nM* TRAS, 2.290 *eV* and 0.118 *eV* for 100 *nM* TRAS, 2.287 *eV* and 0.116 *eV* for 200 *nM* TRAS respectively. This tells us that the very small fraction of HER2 causes the redistribution of the perturbation potential equivalent to the strongest antibody-antibody interaction observed for concentration around 200 *nM*.

As can be seen from Fig. 5a) and b), increasing the HER2 concentration in the samples containing the three above-stated concentrations of the TRAS leads to an increase in the emission energy and emission width. This can be considered as the effective perturbation potential recovering caused by the increasing HER2 concentration. Thus, at HER2-HER2 distance of 140 *nm*, corresponding to 600 *nM* of HER2, the emission energy and emission width for three TRAS concentrations are almost equivalent to the values measured for the corresponding concentrations of the TRAS antibodies only. This allows us to conclude that reduction of TRAS-HER2 interaction is caused by increasing HER2 concentration.

The dependencies of the TRAS-HER2 emission energy and emission width measured for the samples with 30 *nM*, 100 *nM*,and 200 *nM* TRAS were modeled with Eq. 1.1 and Eq. 1.2. The antibody-antigen interaction-induced emission 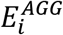 in this case appears to be positive and decreases with increasing initial TRAS concentration16 *meV*, 4.5 *meV*, and 1.4 *meV*, respectively. Similarly, the antibody-antigen interaction-induced emission broadening 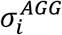 decreases with increasing antibody concentration 10 *meV*, 3 *meV*, and 2 *meV*, respectively. The parameter *n*= 4 reflects the complexity of the TRAS-HER2 interactions and governs transition between interaction induced narrowing and broadening. The interaction transfer length 180 *nm* and 120 *nm* for the emission energy and for the emission width, and the interaction transfer parameter *β* also appears to decrease for higher TRAS 1, 0.5, and 0.25, respectively.

The final experiment consisted of measurements of the TRAS-HER2 interaction in the presence of serum at the same dilutions discussed above, namely 1: 50, 1: 25, 1: 10, and 1: 0. TRAS samples with concentrations of 30 *nM*, 100 *nM*, and 200 *nM* were investigated at each serum dilution with increasing HER2 concentration. All measured dependences exhibit qualitatively similar behavior, see Fig.6. A small initial increase, followed by a comparable decrease in shorter distances. The fitted value of interaction-induced emission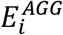, for increasing HER2 concentration, is small, around 1 *meV*. The peak of the dependence is shifted toward longer distances, indicating that the interaction transfer length is longer than for the samples measured in pure HBS-8, serum suppresses strong TRAS-HER2 effect.

Under excitation condition of the experimental setup [18,19] undiluted serum exhibits emission overlapping with the emission of Rh6G. Numerical subtraction of the signal originating from serum distorted the emission signal measured for the sample containing 30 *nM* TRAS and resulted in lower signal to noise (SNR) ratio. This resulted in some dispersion of the emission energy and significant dispersion of the emission width, see Fig. 6g) and h) with open red squares. In all other TRAS concentrations 100 *nM* and 200 *nM*, the fluorescence was sufficiently strong to achieve a good SNR and allowed us to carry on reliable analysis.

**Fig. 6.**
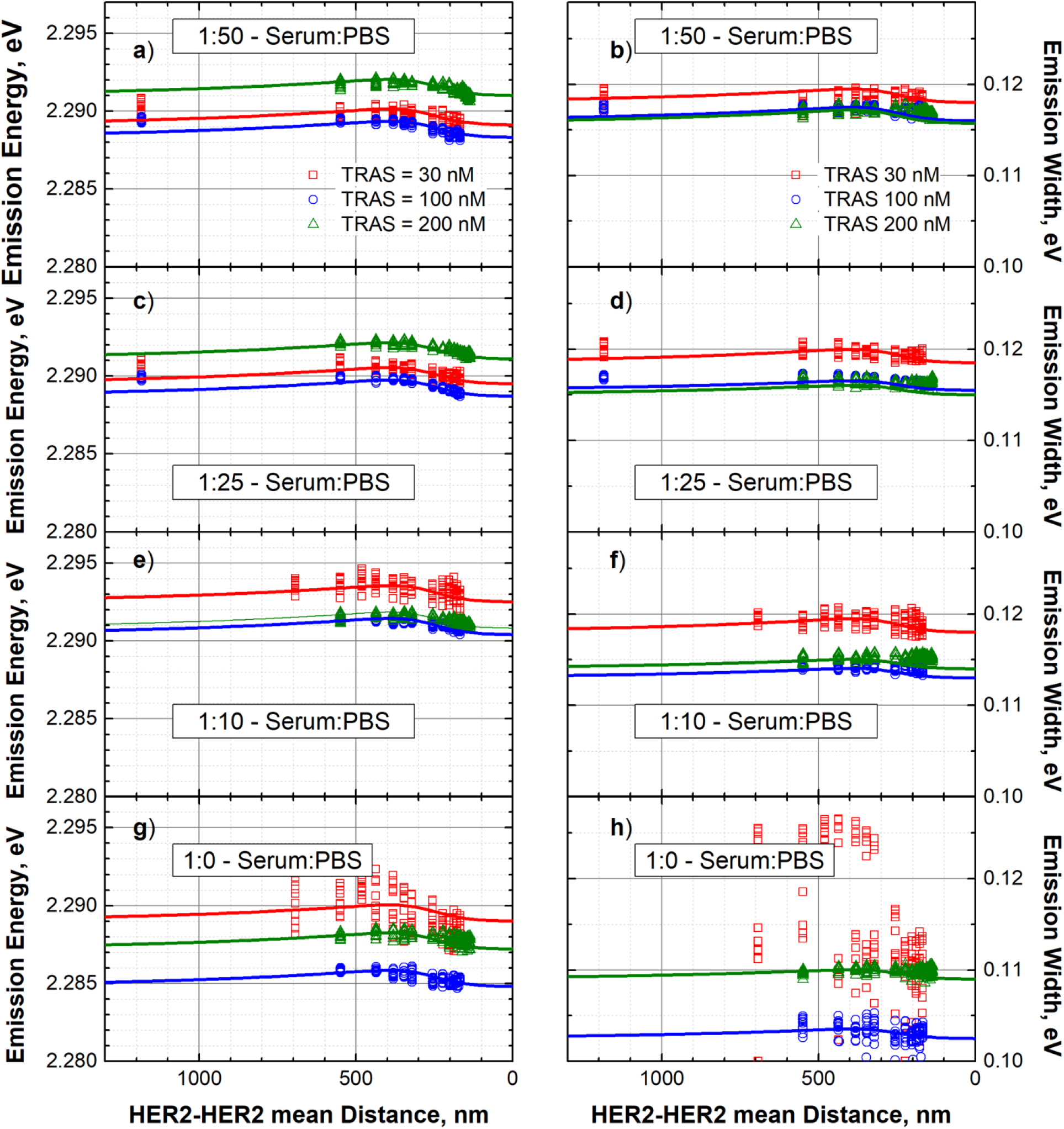
Dependence of emission energy **a), c), e), g)** and emission width **b), d), f), h)** of dye conjugated TRAS at three initial concentrations **30 *nM*** (red squares), **100 *nM*** (blue points) and **200 *nM*** (green triangles) on HER2-HER2 mean distance (concentration) measured in **1: 50, 1: 25, 1: 10 and 1: 0 serum**serum. Experimental data are shown with the points, theoretical models with solid lines of respective color.

The parameter that obviously depends on increasing serum content is the constant offset 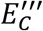. However, as seen in Fig. 6, it also depends on the initial concentration of TRAS. The emission width also changes with increasing serum content. Serum dilutions 1: 50 and 1: 25 show a remarkable difference in the TRAS-HER2 dependence between the low TRAS concentration of 30 *nM* (~400 *nm*) and the higher TRAS concentrations of 100 *nM* (~255 *nm*) and 200 *nM* (~200 *nm*), where the higher concentrations produce a narrower emission width.

From Fig. 6 it is noticeable that increasing HER2 concentration has a minor effect on the emission energy of dye-conjugated TRAS. It remains unclear whether TRAS-TRAS interactions contribute to the observed spectral response to the same extent as in pure HBS-8, although it is reasonable to assume that TRAS-TRAS-interaction induced emission shift can be considered the ground state for estimating the strength of the TRAS-HER2 shift of the emission energy. Next, it is remarkable that serum dilution has a major impact on the emission energy. Fig. 7 collects all determined TRAS-HER2 emission energies as a function of serum protein-protein mean distance for 30 *nM*, 100 *nM*, and 200 *nM* dye-conjugated TRAS. All samples show a similar trend, at first the emission energy increases with decreasing serum protein-protein mean distance, and for undiluted serum a substantial drop in the emission energy is observed for all TRAS concentrations. The strongest drop in undiluted serum is observed for the sample with 100 *nM* TRAS.

**Fig. 7.**
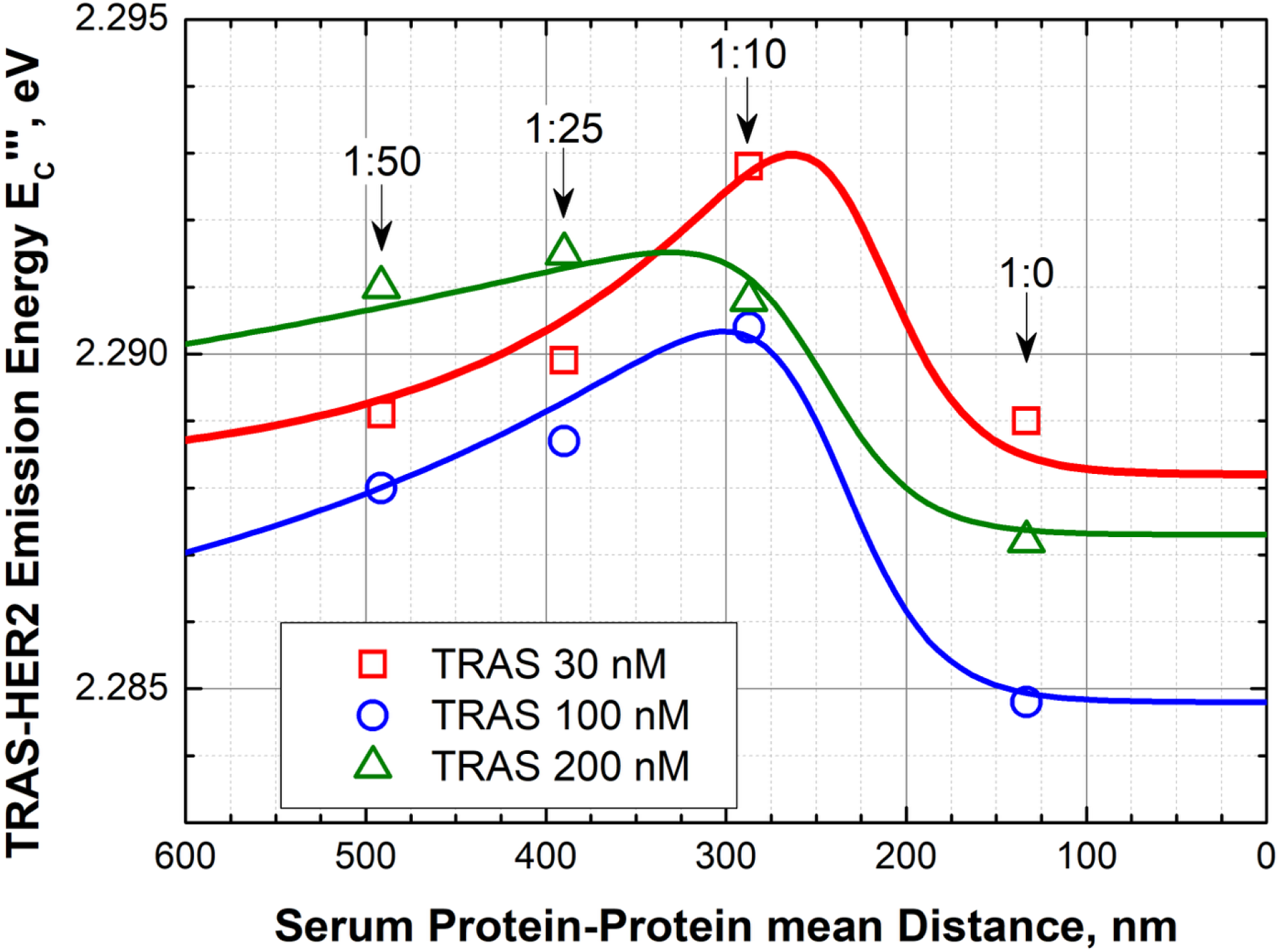
Dependence of TRAS-HER2 emission energy on serum dilution is shown as a function of mean protein-protein distance. The dependencies were obtained for three initial concentrations of the TRAS antibody **30 *nM*** (red squares), **100 *nM*** (blue points), and **200 *nM*** (green triangles). Solid lines with respective colors show theoretically modeled dependencies.

The dependencies shown in Fig. 7 were modeled with Eq. 1.1 and are shown with solid lines. The modeled dependencies have the same value for the parameter *n*, i.e., *n*= 6. The interaction transfer distance is in the range of λ = 140 − 155 *nm*, and *κ* = 5 where decreasing interaction transfer parameter *β* = 1, 0.5, and 0.25 for 30 *nM*, 100 *nM*, and 200 *nM* TRAS, respectively. The interaction-induced emission also decays with increasing TRAS concentration 7.5 *meV*, 3.3 *meV*, and 1.5 *meV*, respectively. It must be noted that the modeled dependencies reflect tendencies and at this stage cannot be treated as absolutely accurate, but some discrete points of serum dilution can be reliably used to determine the TRAS-HER2 interaction-induced change of the emission energy, especially the values measured for undiluted serum. From Fig. 2 i), the TRAS-TRAS emission energies in undiluted serum for 30 *nM*, 100 *nM*, and 200 *nM* TRAS are around 2.292 *eV*, 2.290 *eV*, and 2.289 *eV*, respectively. The TRAS-HER2 emission energies according to Fig. 6 for the same TRAS concentrations are 2.2882 *eV*, 2.2848 *eV*, and 2.2873 *eV*. This estimates the emission energy reductions of 3.8 *meV*, 5.2 *meV*, and 1.6 *meV*, respectively, are likely caused by the TRAS-HER2 interaction. The largest emission energy reduction was observed for the TRAS sample with 100 *nM* concentration indicates that this range of TRAS concentrations is preferable for HER2 proteins to interact with the antibody.

### Conclusions

In this work, the interaction of the dye conjugated TRAS antibody with a recombinant HER2 protein was systematically investigated. The effective perturbation potential acting on dyes conjugated to antibodies such as TRAS and PERT was evaluated through changes in fluorescence emission energy and emission width. These parameters were evaluated from the fluorescence spectra and were used to characterize intermolecular interactions in HBS-8, in diluted and undiluted human serum.

The results demonstrate that antibody-antibody interactions generate measurable changes in the emission energy and the emission width of the dye conjugated TRAS and PERT. Furthermore, binding of TRAS to HER2 produces distinct spectral changes, allowing the interaction to be resolved through fluorescence measurements. In HBS-8, increasing HER2 concentration leads to an increase in emission energy and emission width, whereas in serum-containing media TRAS-HER2 interaction becomes substantially weaker. The magnitude of changes in the emission energy and the emission width as a result of TRAS-HER2 interaction is primarily defined by the concentration of TRAS. The largest interaction-induced emission-energy shift was observed for 100 *nM* TRAS.

The analysis further indicates that proteins present in serum are participating in the transfer of intermolecular interactions, reducing distance-dependent decay and extending the effective range of the interactions. Presented results provide new insight into antibody-antibody and antibody-antigen interactions in complex biological media. They demonstrate that fluorescence spectroscopy can be used to quantify interaction-induced energy changes. In addition, these findings may also have practical applications in the optimization of antibody formulations and interaction-based bioassays.

## Author Contributions

Conceptualization and Investigation: Dinesh Thiyagaraj, Oleh Fedorych.

Methodology, Formal Analysis, Visualization: Oleh Fedorych.

Writing – original draft: Oleh Fedorych.

Writing – review & editing: Dinesh Thiyagaraj, Oleh Fedorych, Alessandro Del Re.

Resources – Duc Quan Pham, Irene Gómez Garrote, Ahmad Saudi

## Supporting Information

The Hamiltonian of the conjugated system is formulated as:

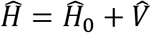

where 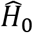 is zero-order Hamiltonian resulting in the mean emission energy of the dye *E*_*D*_, and covalent conjugation and the related local microenvironment perturbations to the electronic states are described by the perturbation potential operator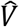.

Applying first-order time-independent perturbation theory, the transition energy of a single dye conjugated to a molecule, *E*_*C*_, can be expressed as:

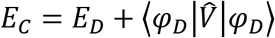

The shift in the mean emission energy, Δ*E*, corresponds directly to the ensemble expectation value of the perturbation potential:

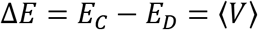

For an ensemble of conjugated dyes,

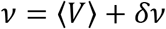

where ∂ν represents the local potential fluctuation from the ensemble average, and ⟨δν⟩ = 0. The mean-square perturbation potential

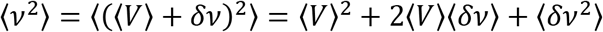

simplifies to

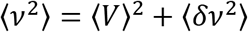

where the ensemble variance ⟨δν^2^⟩ governs the inhomogeneous broadening of the fluorescence spectrum:

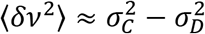

where σ_*C*_ and σ_*D*_ - are the emission widths of the conjugated dye and the free dye, respectively.

The effective perturbation estimated from emission energy and emission width of free and conjugated dye can be written as:

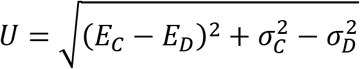

