## Supplementary for "Influence of antibody-antigen interaction on fluorescence of Rhodamine 6G conjugated to the antibodies measured in human serum"

[1]ICHORtec GmbH; Berlin, Germany
*Corresponding authors:

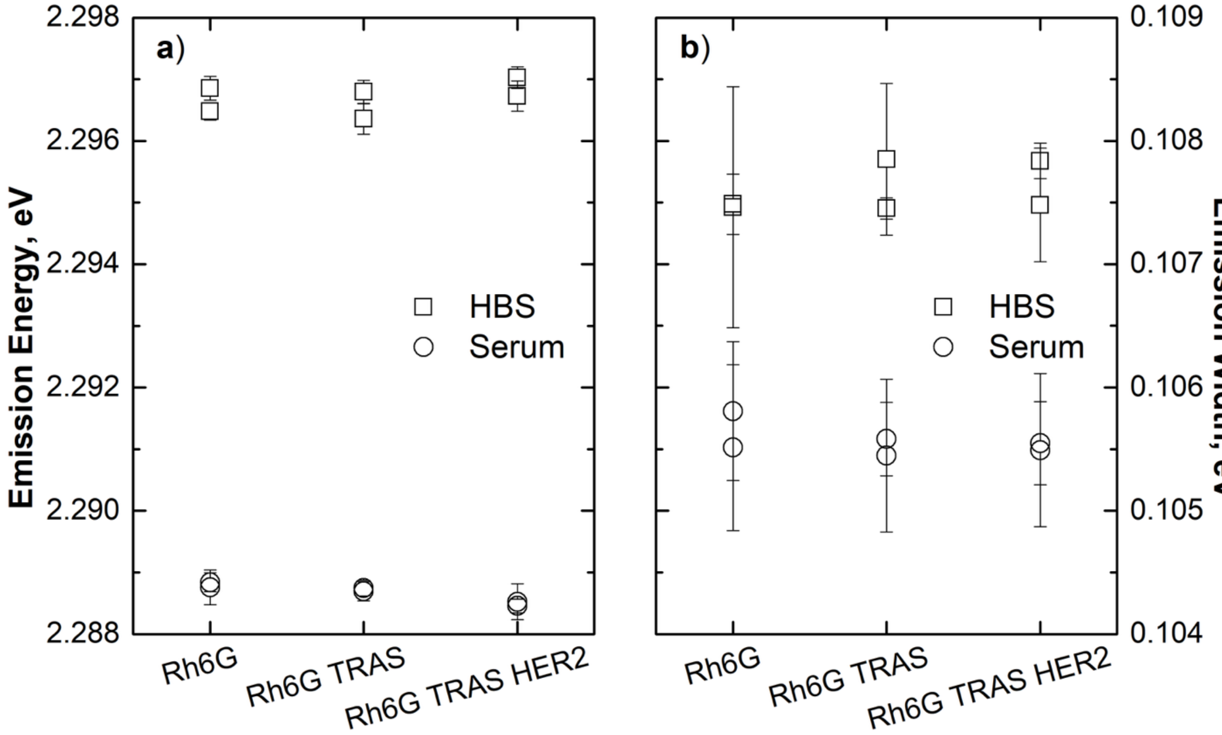


**S1 Fig.** Emission energy a) and emission width b) for $\mathbf{100}\ \boldsymbol{nM}$ of free rhodamine 6G diluted in HBS-8 (squares) and serum (open bullets). Afterwards, in the buffers, $\mathbf{100}\ \boldsymbol{nM}$ of TRAS and $\mathbf{100}\ \boldsymbol{nM}$ of HER2 were added consequentially.

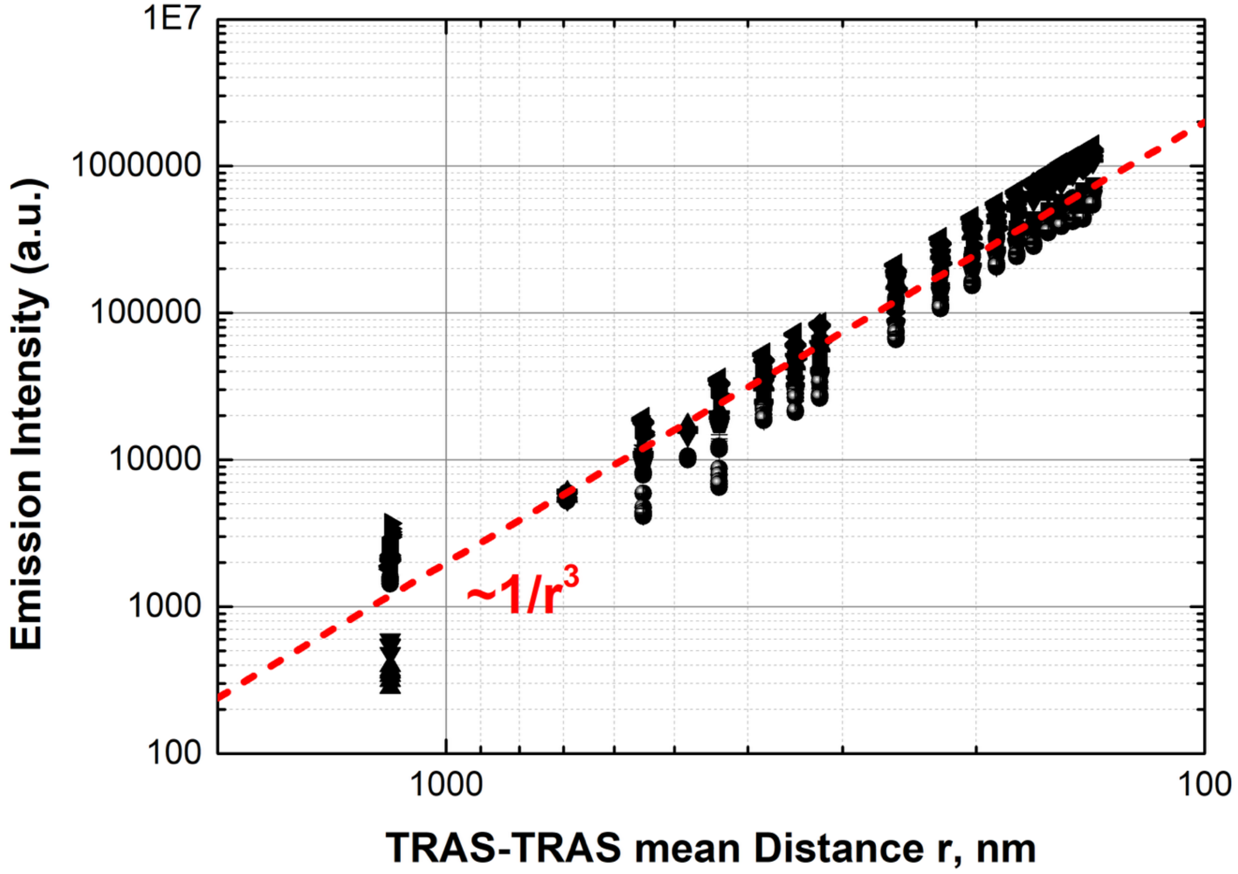


**S2 Fig.** Emission intensity of dye conjugated trastuzumab antibody as a function of it concentration represented in mean distance. Red dashed line shows expected (modeled) increase of the emission intensity on decreasing intermolecular distance (increasing concentration) of TRAS antibody.
